# ASC contributes to sustained neuroinflammation and mild cognitive impairment after closed-head injury

**DOI:** 10.1101/2025.06.19.660521

**Authors:** Tao Li, Sergio Castro-Gomez, Pablo Botella Lucena, Ana Vieira-Saecker, Sarah Schwartz, Yushuang Deng, Yingying Ding, Verena Stein, Eicke Latz, Michael T. Heneka

**Affiliations:** Institute of Physiology II, University Hospital Bonn, University of Bonn, Bonn, Germany; Center for Neurology, Clinic of Parkinson, Sleep and Movement Disorders, University Hospital Bonn, University of Bonn, Bonn, Germany; Luxembourg Centre for Systems Biomedicine (LCSB), University of Luxembourg, Esch-sur-Alzette/Belvaux, Luxembourg; German Center for Neurodegenerative Diseases (DZNE), Bonn, Germany; Institute of Innate Immunity, University Hospital Bonn, Bonn, Germany; Centre of Molecular Inflammation Research, Norwegian University of Science and Technology, Trondheim, Norway; Division of Infectious Diseases and Immunology, University of Massachusetts Medical School, Worcester, USA; Deutsches Rheuma-Forschungszentrum (DRFZ), Berlin, Germany

## Abstract

Mild brain traumatic brain injury (mTBI) from closed-head injuries (CHI) can lead to prevalent neuropsychiatric disorders, including an increased risk for neurodegenerative diseases and dementia. Inflammasomes are molecular complexes crucial for neuroinflammation and secondary damage after trauma, however their role in mild CHI is poorly understood. In this study, we investigate the cellular expression of inflammasome-related genes and their functional significance in CHI models. Single-cell RNA sequencing of cortical tissue revealed selective expression of *Pycard* (PYD and CARD domain containing), also known as *Asc* (apoptosis-associated speck-like protein containing a caspase recruitment domain), which encodes the inflammasome adaptor ASC, predominantly in microglial clusters. Sustained upregulation of inflammasome-related proteins persisted up to 21 days in a model for mTBI, with this pattern significantly reduced in *Asc^−/−^* mice. Importantly, mild cognitive impairment induced after mild CHI was largely abrogated in *Asc^−/−^* mice. These findings suggest that ASC, as the primary inflammasome adaptor, plays a critical role in sustaining neuroinflammation and contributes to cognitive deficits after mild CHI. This study provides insights into the molecular neuroinflammatory mechanisms underlying CHI, potentially informing future therapeutic strategies.

## Introduction

Traumatic brain Injury (TBI) represents a significant global health challenge and is considered a silent epidemic, affecting individuals across all demographics and age groups. Every year, TBI affects around 1.5 million people in both the European Union and the United States. In the EU, these injuries are linked to about 57,000 deaths and 1.5 million hospital admissions (1), while in the U.S. they cause roughly 50,000 deaths, 230,000 hospitalizations with survival, and 80,000–90,000 new cases of long-term disability (2). The majority of cases are clinically classified as mild TBI (mTBI), typically resulting from closed-head injuries (CHI) caused by falls, traffic accidents, violence, contact sports, or military actions (3, 4). Although mTBI symptoms can recover within days or weeks, up to 20% of individuals experience persistent physical, cognitive, and behavioral impairments that lead to a reduced quality of daily life and an elevated risk of neuropsychiatric disorders, including depression and dementia (5–7). Despite its prevalence, the underlying pathophysiology of mTBI resulted from CHI remains poorly understood, posing great challenges for early clinical diagnosis and timely intervention.

Reactive microglia and astrocytes play crucial roles in the innate immune response and secondary damage following TBI. Prolonged activation of these cells impairs debris clearance, contributing to neuroinflammation, exacerbating neuronal dysfunction and damage, and promoting abnormal protein aggregation (8–10). Advances in imaging techniques have allowed the *in vivo* visualization of glial reactivity after CHI, emphasizing the involvement of microglia and astrocytes in the inflammatory cascade that contributes to the development of post-traumatic disorders (11–13). A key element in initiating and maintaining innate immune responses of the brain is inflammasome activation and signaling. Inflammasomes are oligomeric protein scaffolds rapidly activated by some damage-associated molecular patters (DAMPs), and organized as a tripartite complex: a cytosolic danger sensor such as NOD-like receptor family pyrin domain containing 3 (NLRP3), a unique adaptor protein called apoptosis-associated speck-like protein containing a CARD domain (ASC), and a proteolytic effector caspase-1 (CASP1) (14). ASC, an intracellular adaptor protein common to almost all inflammasomes, oligomerizes into highly cross-linked macromolecular assemblies, forming visible ASC aggregates. The fibrillar ASC acts as a molecular platform to recruit pro-CASP1, which optimizes signal transduction via proximity-induced CASP1 activation. Activated CASP1 catalyzes the maturation of pro-forms of Interleukin-1β (IL-1β), IL-18 and mediates the proteolytic activation of Gasdermin D (GSDMD). Oligomerized GSDMD forms pores in the plasma membrane, allowing the release of IL-1β, IL-18 and other proinflammatory cytokines. The detection of tissue ASC aggregates is considered a typical hallmark of inflammasome activation (15).

In humans, some inflammasome proteins are detected in blood and cerebrospinal fluid after TBI, and circulating levels of some proteins such as ASC and CASP1 have been proposed as biomarkers for determining injury severity (16–18). Furthermore, inflammasome activators, including NLRP1 (19), NLRP3 (20, 21), and absent in melanoma 2 (AIM2) (22), presumably contribute to the innate immune response and functional disability in mouse models of TBI. these studies primarily focused on inflammasome activation in models of moderate to severe TBI, such as the commonly used Controlled Cortical Impact (CCI), in which ASC has been hypothesized to be dispensable (23) . At the same time, their contribution to mild CHI remains largely unexplored. In the present study, we hypothesized that ASC inflammasomes ensembles contribute actively to the secondary damage following CHI. We aimed to investigate their role in neuroinflammatory responses and subsequent cognitive functions by analyzing a well-established single-cell RNA sequencing (scRNA-seq) dataset of cortical cells from mice subjected to a diffuse TBI model (26) and by using an additional CHI mouse model that induces mild cognitive impairment (27). Our scRNA-seq data analysis reveals a subacute expression of *Asc* (also known as *Pycard*) in cortical cells from mice subjected to midline fluid percussion injury (mFPI), particularly in microglia subpopulation. In concordance with this analysis, we observed sustained upregulation of inflammasome-related proteins, including NLRP3, ASC, CASP-1, and IL-1β, following CHI, persisting up to 21-days post injury (dpi). This expression pattern was significantly reduced in *Asc^−/−^* mice. Moreover, *Asc ^−/−^* mice were protected from the mild cognitive impairment induced by our CHI model, as assessed by the Morris Water Maze (MWM). We also observe ASC-dependent microglial and astrocytic reactivity and interactions that persists over time, accompanied by the upregulation and aggregation of the ASC protein. Our findings identify ASC, the primary inflammasome adaptor, as a key molecule in sustaining neuroinflammation and contributing to cognitive deficits after CHI.

## Results

### Subacute Asc (Pycard) expression in microglia subpopulations after a model for CHI

To examine the cell expression of gene coding for inflammasome-related proteins after CHI, our first approach was to adopt a single-cell genomic analysis. We chose to re-examine a publicly available scRNA-seq dataset from dissociated cortical cells 7 days (7 days post injury, 7 dpi) followed mFPI (26). mFPI mimics several relevant features of CHI including diffused trauma without localized lesion or hemorrhages, and closely models mild to moderate TBI (28). After quality control, we analyzed 14688 control cells and 10,745 cells in the mFPI group (Figure 1A and supplemental Figure 1A). The complex multidimensional dataset was transformed and visualized using the nonlinear dimensionality reduction technique Uniform Manifold Approximation and Projection (UMAP) (Figure 1A, 1B, 1E). Examination of cellular transcriptional identities at the single-cell level uncovered 22 distinct cellular clusters. Cellular subpopulation labels were assigned via automated cell type annotation per media sample using SingleR (29) (Supplemental Figure 1B). Cells identified as microglia were specially enriched in this dataset (Figure 1B). A differential expression gene analysis of this subset of cells showed a typical disease-associated microglia signature after TBI, with overexpression of type-1 interferon genes (such as *Ifi27l2a, Ifitm3, Ifit3, rf7*) and *Apoe,* and reduction of homeostatic genes such as *tmem119* or *tgfb1* (Figure 1C).

**Figure 1:**
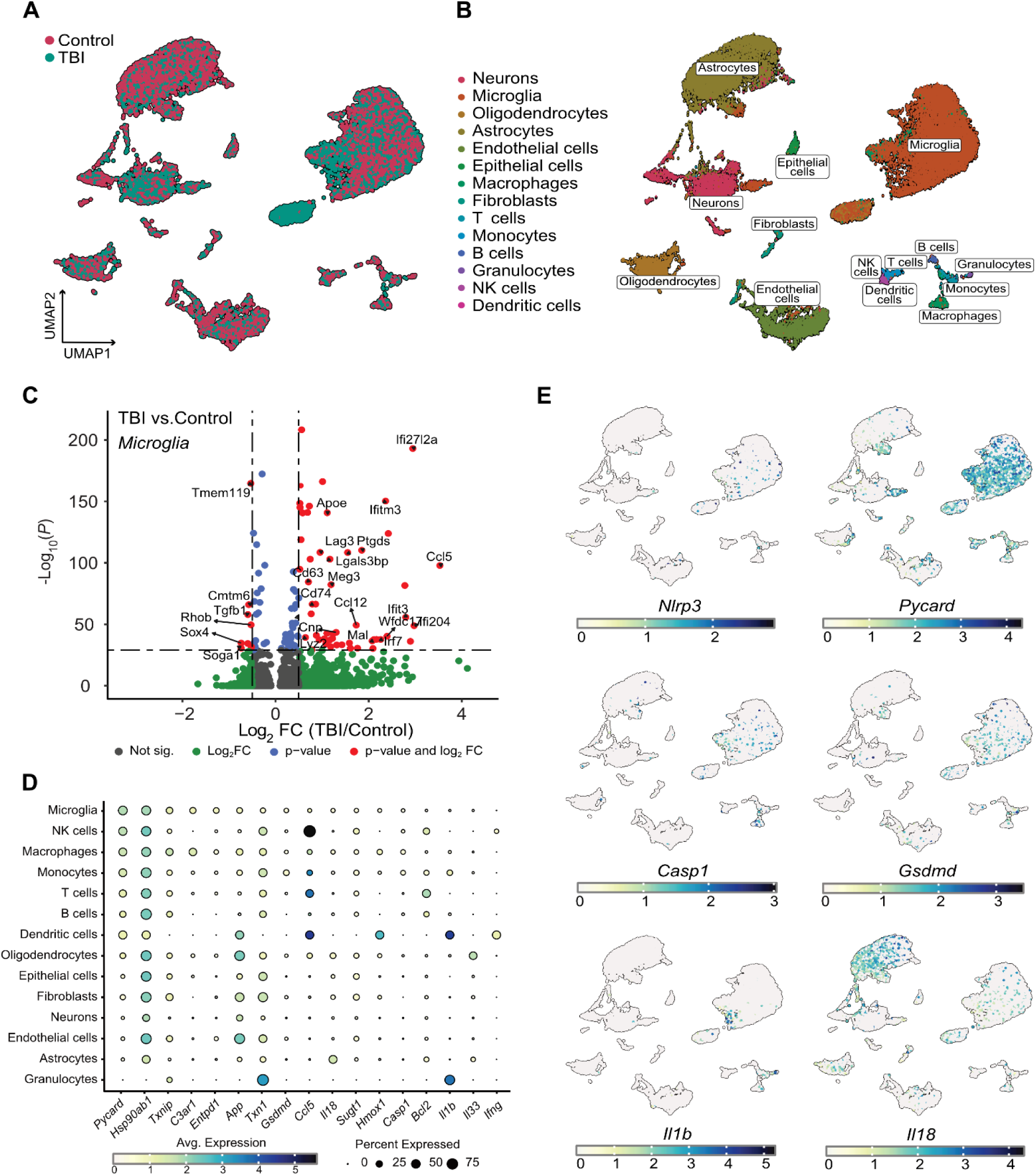
Subacute *Asc (Pycard)* expression in microglia after a model for CHI. A-B. Single-cell RNA sequencing (scRNA-seq) analysis of brain suspended cells from mice 7 days after mFPI. **A.** UMAP visualization showing clustering of different cell populations, with mFPI and control samples overlaid. **B.** Annotated cell types, including microglia, macrophages, and other immune and neural cells after cluster identification. **C.** Volcano plot depicting differentially expressed genes in microglia of control and mFPI samples. **D.** Dot plot expression analysis showing percent of expressed key inflammasome-related genes across various brain cell types, highlighting *Asc* (*Pycard*) expression in microglia, Natural Killers (NK) cells, macrophages and monocytes. **E.** UMAP projection of cells colored relative to the expression of canonical inflammasome-related genes *Nlrp3*, *Asc* (*Pycard*) *Casp1, Il1b, Il18 and Gsdmd*.

We then evaluated, for each predicted cellular subpopulation, the expression of genes annotated in pathways related to inflammasome activation (pathcards.genecards.org (30), Supplemental Fig 1C). Notably, we found *Asc* (or *pycard,* gene coding for the adaptor protein ASC), as one of the genes with wider expression in clusters represented by microglia, macrophage, and other peripheral infiltrating immune cells (Figure 1D-E and Supplemental Figure 1C, 1D). These clusters show a more reduced percentage of cells expression genes coding for inflammasome effectors such *Gsdmd, Casp1, il1b* or *il18* (Figure 1D and E). Surprisingly, we found a very low percentage of cells expressing genes coding for inflammasome activators such as *Nlrc4, Nlrc5, Nlrp1b, Nlrp3 or Aim2* (Supplemental Fig 1C). These results suggest that the subacute and prolonged expression of the adaptor protein ASC, especially in microglia, may play a central role in the sustained innate immune response and pathophysiology of TBI related to CHI.

### ASC contributes to the sustained expression and processing of inflammasome mediators following mild CHI

To unveil the pathophysiological role of inflammasomes and specifically of ASC as main inflammasome adaptor and abundantly expressed gene in microglia after CHI, we adopted an electromagnetic CHI procedure as main model for mTBI (31). In this model, a controlled cortical impact injury is delivered directly to the midline surface of the skull using a 5 mm diameter tip at a specified depth and velocity (Supplemental Figure 2A). We found mild motor, reflex or reaction deficits at 1- or 7 dpi when the mice were evaluated using the Revised Neurobehavioral Severity Scale (NSS-R) for rodents (32) (Supplemental Figure 2B). To examine the protein expression of inflammasome-related proteins after CHI, we performed immunoblot analysis of cortices from WT and *Asc^−/−^* mice that underwent either sham or CHI surgery. We analyzed and semi-quantified the protein levels of NLRP3, ASC, CASP1, and IL-1β at 1, 7, and 21 dpi (Figure 2A-I). Our results revealed sustained upregulation of these proteins up to 21 days after CHI in WT mice. Notably, mice with a constitutive genetic deficiency of the *pycard* gene (*Asc^−/−^*) showed a very significant deficit in the expression and cleavage of CASP1 (Figure 2D-E) and of IL-1β over time (Figure 2G) fallowing CHI. ELISA measurements further demonstrated significantly lower soluble levels of IL-1β and TNF-α in *Asc^−/−^* mice (Figure 2H-I), particularly at later time points. We additionally evaluated the levels of cleavage of Caspase-8 (CASP8), a proposed non-canonical enzyme involved in IL-1β processing (33) (Supplemental Figure 2C and D), and Caspase-3 (CASP3), as indirect measurement of apoptotic pathways (34) (Supplemental Figure 2C and E). Consisting with the reduction in soluble IL-1β, *Asc^−/−^* cortices showed significantly reduced levels of cleaved CASP8 up to 21 days post CHI (Supplemental Figure 2D), which suggest parallel activation of canonical and non-canonical pathways of IL-1β processing, both of which appear largely dependent on ASC. Additionally, the cleavage of CASP3 was significantly reduced in *Asc^−/−^* mice (Supplemental Figure 2E), suggesting a strong neuroprotective effect and a complex crosstalk involving the activation of multiple types of cell death influenced by inflammasome signaling (35).

**Figure 2:**
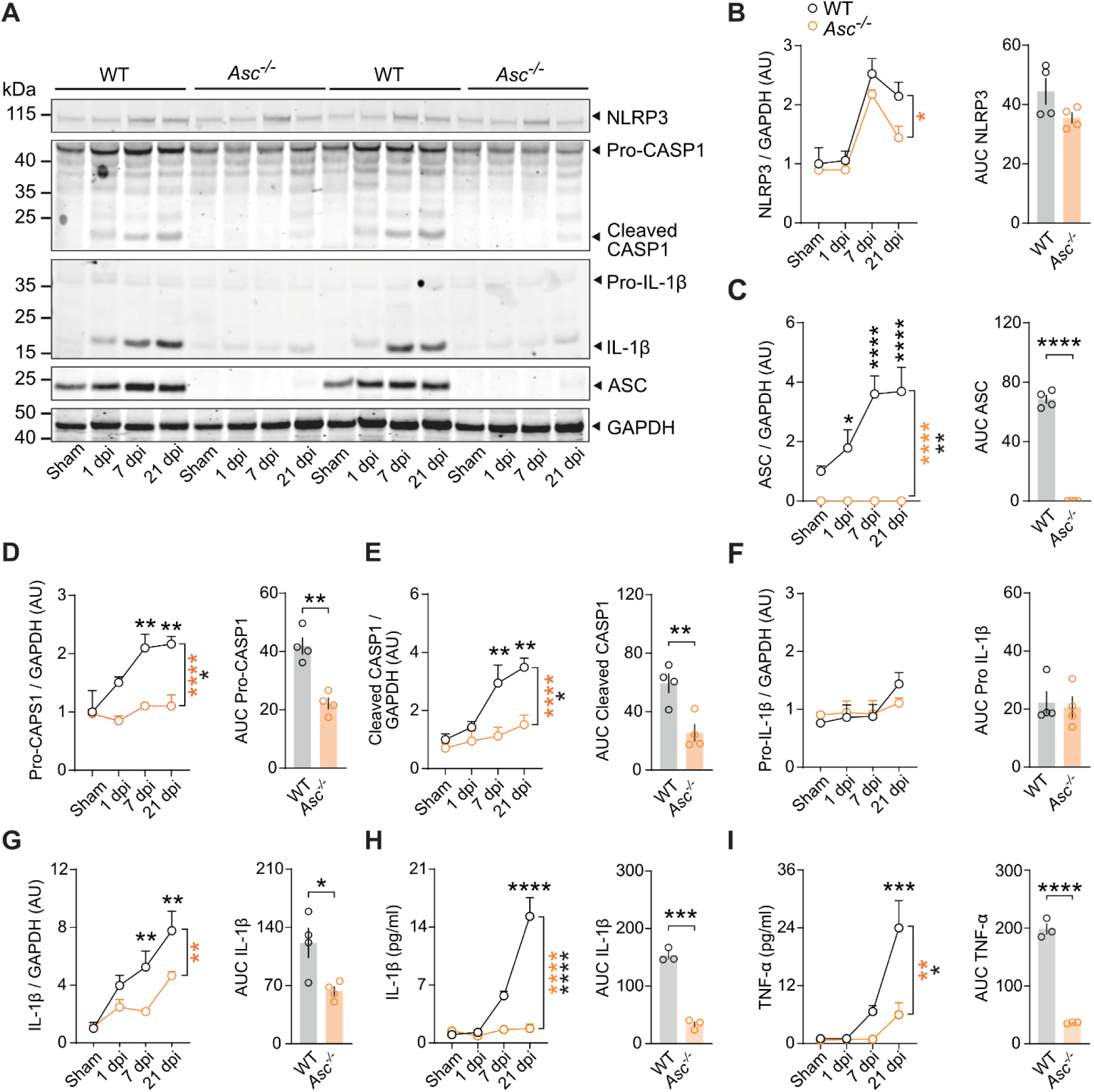
ASC contributes to the upregulation of inflammasome mediators following CHI. **A.** Immunoblot detection and **B-G.** densitometric semi-quantification of inflammasome-related proteins NLRP3, ASC, Caspase-1 (CASP1), IL-1β at 1, 7, and 21 dpi in contusional cortices from WT and *Asc^−/−^* mice following Sham or CHI. (left panels *n* = 4 per group and time point, \**p* < 0.05, \*\**p* < 0.01, \*\*\**p* < 0.001, \*\*\*\**p* < 0.0001, Ordinary two-way ANOVA, Bonferroni’s test. AU, arbitrary units. **H, I.** Levels of IL-1β and TNF-α in brain lysates from WT and *Asc^−/−^* mice at 1, 7, and 21 dpi following CHI. *n* = 3 - 4 per group and time point, Left panels represent ordinary two-way ANOVA, Bonferroni’s test, \**p* < 0.05, \*\**p* < 0.01, \*\*\**p* < 0.001. Right panels in **B-I** represent integrated relative concentrations (Area Under Curve, AUC), \*\**p* < 0.01, \*\*\**p* < 0.001 by Student’s t-test. dpi, days post-injury.

To identify the cellular sources of IL-1β following mild CHI, we conducted immunohistochemical staining for IL-1β, ionized calcium binding adaptor molecule 1 (Iba1) and Glial fibrillary acidic protein (GFAP) on cortices of WT mice in sham, 7 and 21 dpi groups (Supplemental Figure 3a-d). At 7 days post-mTBI, IL-1β signal was found to be colocalized with or in the vicinity of GFAP^+^ cells (presumably reactive astrocytes) and with Iba1^+^ cells (likely microglia) (Supplemental Figure 1A-D). Interestingly, at 21 dpi, the majority of IL-1β was detected in Iba1^+^ cells (microglia or infiltrating macrophages) suggesting a very active role of microglia on the TBI-dependent upregulation and secretion of this cytokine (Supplemental Figure 3C-D). Our findings thus emphasize the crucial role of inflammasome signaling, with ASC as the common adaptor protein, in sustaining the expression and processing of neuroinflammatory mediator after CHI. Moreover, we identified Iba1^+^ cells, likely microglia, or infiltrating macrophages, as the primary cellular sources of IL-1β, especially at later time points post-injury.

### ASC shapes the morphology of Iba1^+^ cells following CHI

Microglia exhibit distinct adaptive morphologies in response to mechanical stimuli, and their morphology is considered a proxy for their function (36). To investigate microglia architecture after CHI, we performed immunofluorescence staining for Iba1 and performed skeleton analysis (37) to examine cell morphology in cortices of WT and *Asc^−/−^*mice at 1, 7, and 21 dpi (Figure 3A–B). First, we observed that *Asc^−/−^* mice consistently exhibited lower Iba1+ cell counts compared with WT mice at 7 dpi (Figure 3C), suggesting a decreased microgliosis in the subacute CHI phase. Next, we quantified the normalized changes in the morphology of Iba1+ cells based on cell count. The morphological alterations, including the number of branches per cell, endpoints per cell, and branch length per cell, significantly decreased after injury in WT mice, particularly at 21 dpi. In contrast, *Asc^−/−^* mice showed a significant preservation of these morphological features at 21 dpi compared to WT mice (Figure 3D-F). In conclusion, our observations suggest a wave of microgliosis in the subacute phase and revealed a progressive debranching of Iba1+ cells over time following injury. Notably, the lack of ASC demonstrated protection against these sustained cellular morphological alterations, particularly at later time points.

**Figure 3:**
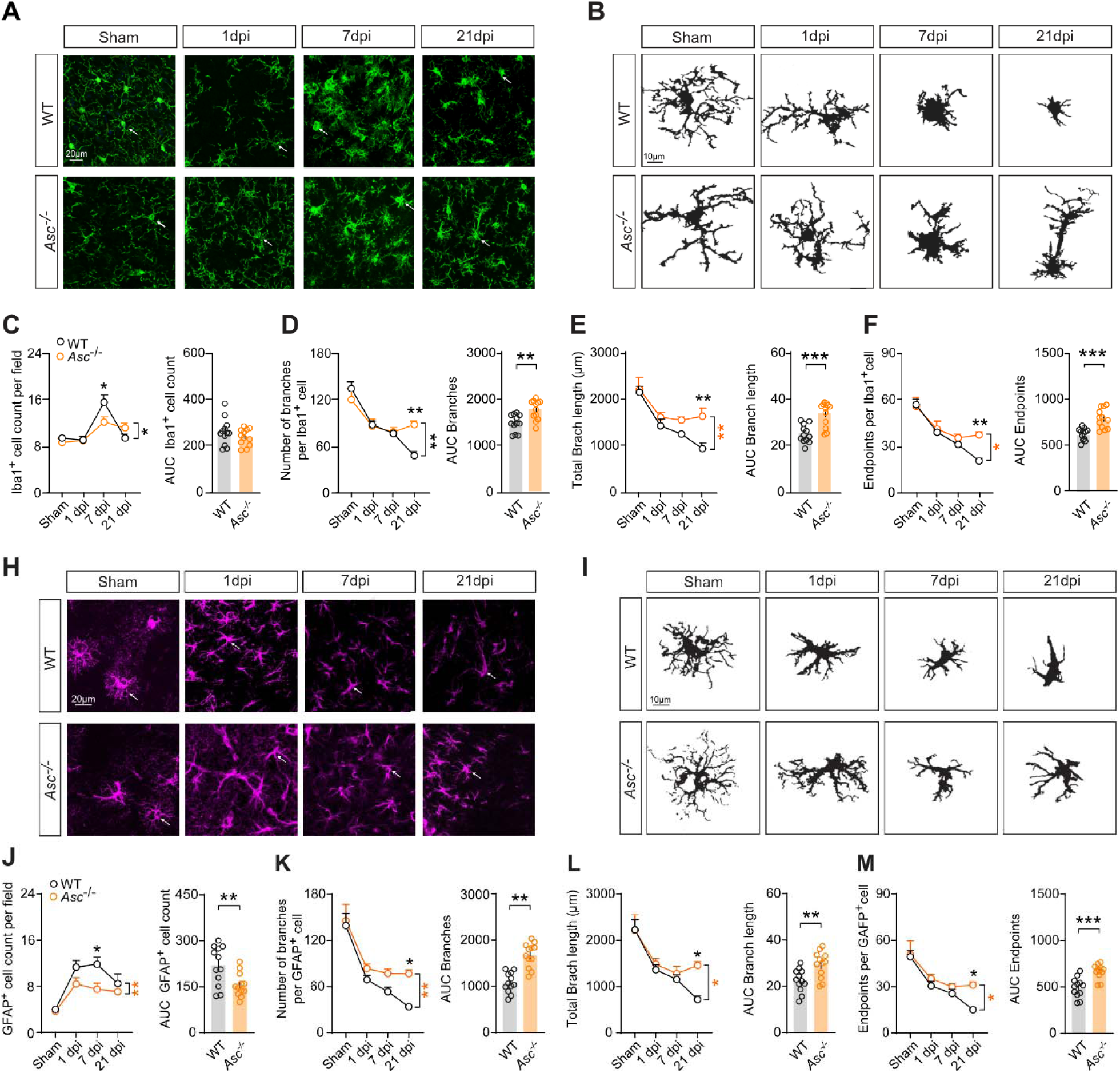
Morphology of Iba1^+^ and GFAP^+^ cells are modulated by ASC following CHI. **A.** Immunohistochemistry of Iba1 (green, presumably microglia) in cortices from WT, and *Asc^−/−^* mice at 1, 7, and 21 dpi following CHI. Arrows indicate representative morphological changes. Scale bar, 20 μm. **B.** Representative skeletonized Iba1^+^ cells following CHI. Scale bars, 10 μm. Quantitative analysis of **C.** Iba1^+^ cell counts; **D.** branch numbers per cell, **E.** total branch length per cell and **F.** endpoint numbers per cell at 1, 7, and 21 dpi in WT and *Asc^−/−^* mice subjected to sham or CHI. **H.** Immunohistochemistry of GFAP (magenta, presumably astrocytes) in cortices from WT and *Asc^−/−^*mice at 1, 7, and 21 dpi following CHI. Arrows indicate representative morphological changes. Scale bar, 20 μm. **I.** Representative skeletonized GFAP^+^ cells following CHI. Scale bars, 10 μm. Quantitative analysis of **J.** GFAP^+^ cell counts; **K.** branch numbers per cell, **L.** total branch length per cell and **M.** endpoint numbers per cell at 1, 7, and 21 dpi in WT and *Asc^−/−^* mice subjected to sham or CHI. *n* = 12 slices from 4 mice per group per each time point, \**p* < 0.05, \*\*\**p* < 0.01, \*\*\**p* < 0.001, \*\*\*\**p* < 0.0001 by ordinary two-way ANOVA with Bonferroni’s post hoc test. Right panels show integrative AUC of each marker over time (dpi). \**p* < 0.05, \*\**p* < 0.01, \*\*\**p* < 0.001, \*\*\*\**p* < 0.0001 by Student’s t-test.

### ASC regulates the reactivity of GFAP^+^ cells following CHI

Previous studies have provided evidence that morphological changes in astrocytes are intricately linked to microglia activation during inflammation-induced responses (38). To examine reactive astrocytes and their morphological alterations following mild CHI, we performed a morphological analysis of GFAP^+^ cells in contusional layers 1 and 2/3 of the retrosplenial and primary motor cortices, using the same method as for Iba1+ cells. (Figure 3H-I). We initially observed a noteworthy increase in the count of GFAP^+^ cells at 1, 7 and 21 dpi in WT mice (Figure 3j), confirming increased reactive astrogliosis in our model. *Asc* deficiency exerted a genotype-related effect in the cell count similar to what we observed in Iba1^+^ cells, showing significantly lower numbers, especially at 7 dpi (Figure 3j). Subsequently, we quantified normalized morphological changes in GFAP^+^ cells based on cell counts. The branches per cell, endpoints per cell, and length of branches per cell were significantly decreased after CHI, particularly at 21 dpi. In contrast, *Asc^−/−^* mice show preserved morphological features at 21 dpi compared to WT mice (Figure 3K–M). Interestingly, at 21 dpi, we observed clusters of aggregated GFAP+ cells, indicating spatial reorganization post-injury. Similar morphological changes were evident in both microglia and reactive astrocytes after injury, suggesting a potential correlation or interactions between these two glial cell populations.

### Iba^+^ and GFAP^+^ cell interactions are modulated by ASC following CHI

To investigate the interaction between reactive astrocytes and microglia in the contusional cortex following injury, we performed surface–surface colocalization analysis of the contact areas after 3D reconstruction of confocal images from GFAP- and Iba1-immunostained tissue (Figure 4A). We observed a significant increase in contact area between reactive astrocytes and microglia at 7 dpi, reaching a peak at 21 dpi in WT mice (Figure 4B). Notably, the contact area was significantly reduced at 7 and 21 dpi in *Asc^−/−^*mice (Figure 4B) which may be explained by the deficient microglia activation and reactive astrogliosis observed in our previous analysis. These findings, together with our expression analysis, suggest that inflammasome ensembles and neuroinflammatory processing are essential for sustained microglial and astrocytic responses and cellular interactions after mild CHI.

**Figure 4:**
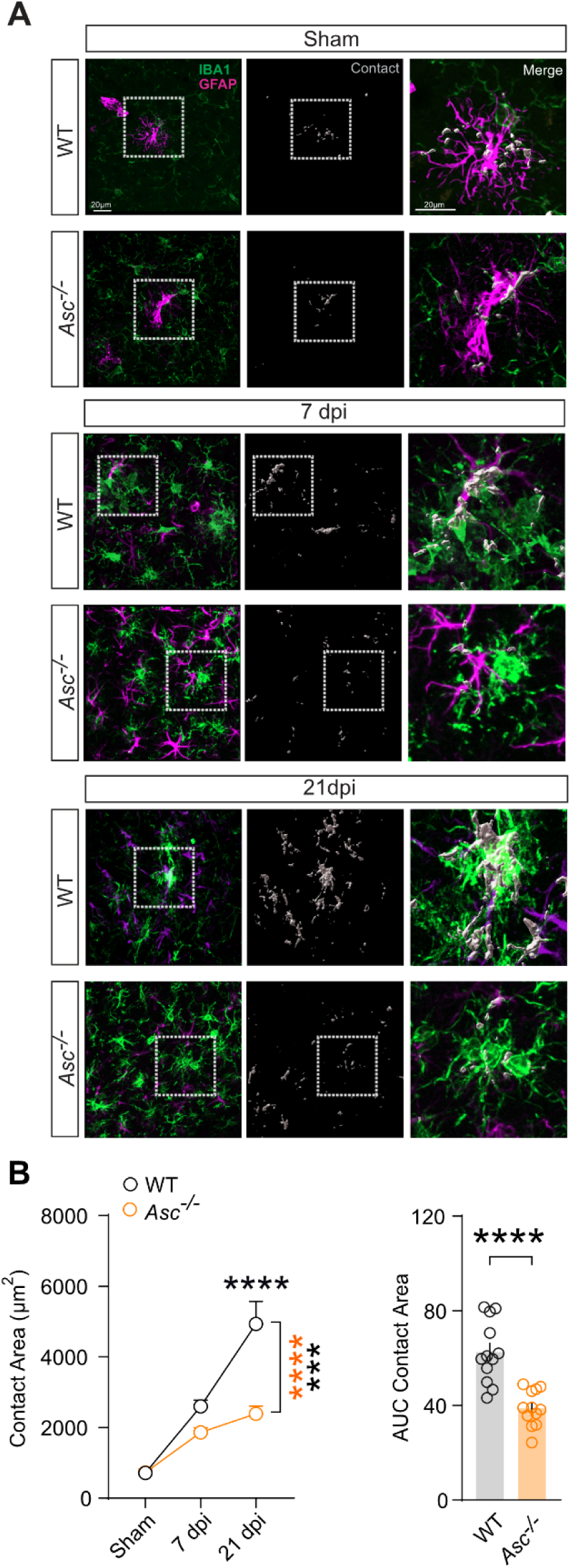
Cellular interactions of microglia and reactive astrocytes are modulated by ASC following CHI. **A.** Immunohistochemistry for Iba1 (green) and GFAP (magenta) in cortices of WT and *Asc^−/−^* mice at 7 and 21 dpi. Surface-surface colocalization analysis after 3D reconstruction was used to estimate contact areas between Iba1^+^ cells and GFAP^+^ cells. Rectangles highlight examples of glial surface-surface colocalization, shown in high magnification views. Scale bar, 20 μm. **B.** Quantification of contact areas between Iba1^+^ cells and GFAP^+^ cells at 7 and 21 dpi in WT, and *Asc^−/−^* mice subjected to Sham or CHI. *n* = 12 slices from 4 mice per group per each time point, \**p* < 0.05, \*\*\**p* < 0.01, \*\*\**p* < 0.001, \*\*\*\**p* < 0.0001 by ordinary two-way ANOVA with Bonferroni’s post hoc test. Right panels show integrative AUC of each marker over time (dpi). \**p* < 0.05, \*\**p* < 0.01, \*\*\**p* < 0.001, \*\*\*\**p* < 0.0001 by Student’s t-test.

### Genetic deficiency in the Asc gene provides protection against mild cognitive impairments following

Having demonstrated that *Asc* is predominantly expressed by microglia in the subacute phase of CHI, contributing to the sustained inflammasome activation, as well as chronic microglial and astrocytic reactivity following CHI, we next sought to investigate whether the genetic removal of the main inflammasome adaptor protein ASC influences cognitive outcomes after mild CHI. In humans, a substantial number of individuals present transient or persistent cognitive impairment following mTBI, including deficits in memory recall, procedural tasks, executive functions, attention and emotional symptoms such as heightened anxiety (39). To investigate how ASC contribute to these neurobehavioral phenotypes, we performed Morris Water Maze (MWM) as main behavioral task in WT and *Asc^−/−^* mice after CHI. MWM was conducted from days 10 to 20 after CHI (10-20 dpi) and consisted in a spatial learning phase (10 to 15 dpi), a 24-hour long-term spatial memory test (16 dpi), and a cued learning phase as control procedure between 17 and 20 dpi (Figure 5A). In comparison to the WT sham group, WT mice subjected to CHI learned the task with comparable daily reduction in path lengths towards the hidden platform (Figure 5B). However, CHI induces significantly prolonged swimming time near the walls (thigmotaxis) during the learning phase in WT injured mice (Figure 5D). We observed similar tendency in thigmotaxis during the cued version of the task (Figure 5H). This procedural deficit was largely reduced in *Asc^−/−^* mice (Figure 5D-E and 5H-I). Additionally, *Asc^−/−^* mice demonstrated preserved long-term spatial memory 24 hours after the learning phase, as visually suggested by the density plots (heat maps) of swim paths (Figure 5J). This finding was further supported by a significantly greater percentage of time spent exploring the target quadrant (Figure 5K), as well as more frequent crossings of the virtual target platform (Figure 5I) compared to the corresponding measures in another quadrant or at other virtual platforms (Figure 5L) during the probe trial performed at 16 dpi. Furthermore, *Asc^−/−^* mice displayed slightly higher velocity in the last day of cued learning (Supplemental Figure 4A and B). These data show that CHI induces mild cognitive impairments, mainly as procedural and memory deficits in the MWM, which are mostly dependent on ASC expression.

**Figure 5:**
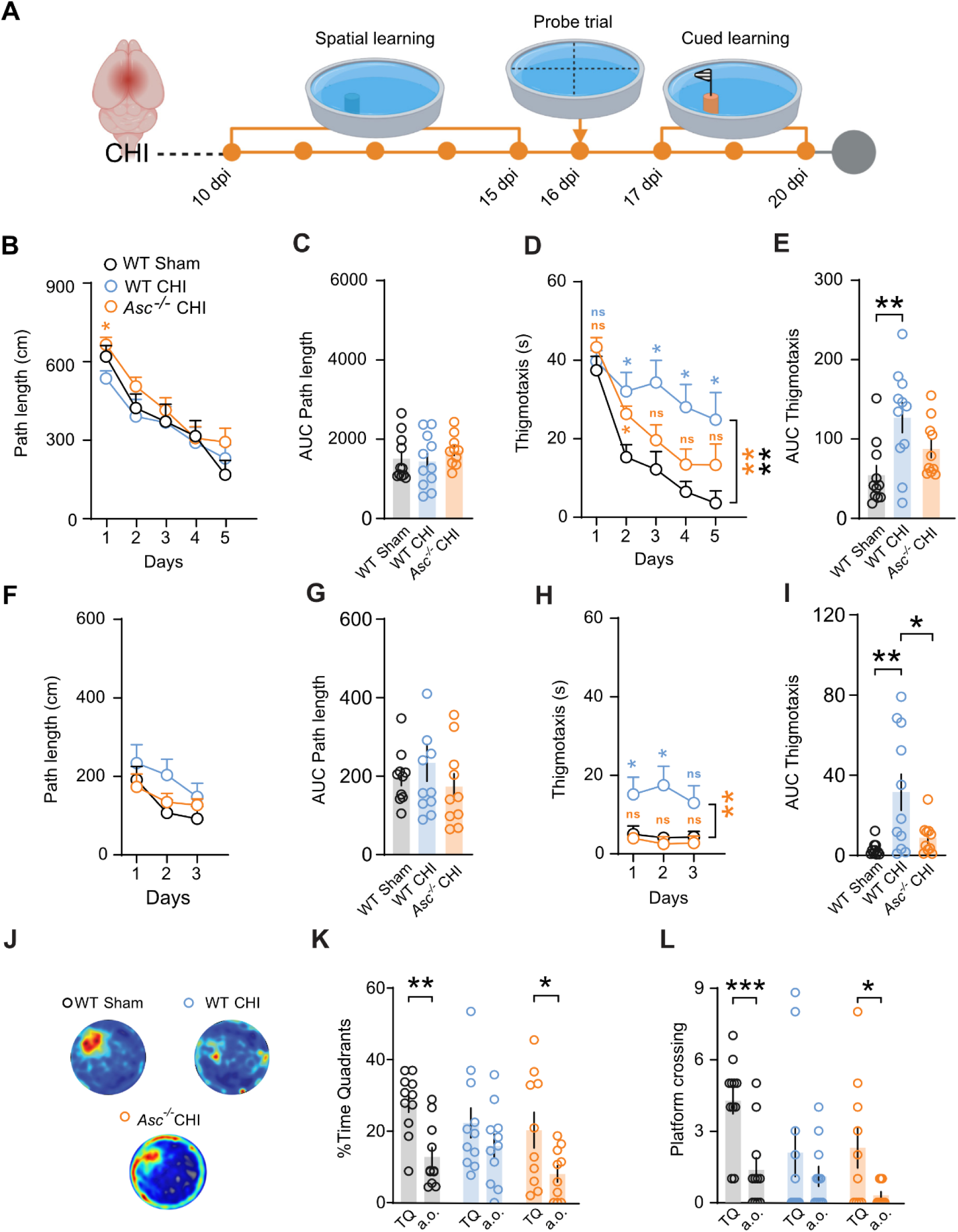
Genetic deficiency of the Asc gene protects against mild cognitive impairments following mild CHI. **A.** Experimental timeline for cognitive assessment following CHI, including spatial learning phase (10-15 dpi), probe trial (16 dpi), and cued learning (17-20 dpi) in the Morris water maze. **B-E.** Morris water maze performance in WT and *Asc*^−/−^ mice, showing path length and thigmotaxic across spatial **B-E** and **F-J** cued training days (n=10-11 mice per genotype *p < 0.05, **p < 0.01, ***p < 0.001, ****p < 0.0001 by repeated measurements two-way ANOVA with Tukey’s post hoc test). Integrative AUC for path length and thigmotaxic from spatial learning (**C** and **E**) and cued learning phase (**G** and **I**). **J.** Representative heatmaps mean swimming trajectories and **k.** quantification of percentage of time spent in target in comparison to another quadrants (a.o.) or **L.** platform crossing times in virtual target platform in comparison to another viral platform (a.o.) (n=10-10 mice per Genotype, *p < 0.05, **p < 0.01, ***p < 0.001 by ordinary one-way and two-way ANOVA with Tukey’s post-hoc tests). Data are presented as mean ± SEM.

### NLRP3 regulates ASC aggregation and distribution following CHI

The NLRP3 is widely recognized as one of the most relevant inflammasome activator in the context of TBI, with growing evidence indicating its central role in mediating neuroinflammation and secondary brain damage (40). To explore the involvement NLRP3 in the aggregation of ASC as the main inflammasome adaptor in post-CHI, we conducted immunohistochemical staining of the contusional cortical tissue for Iba1, GFAP and ASC at 7 and 21 dpi in in WT and *Nlrp3^−/−^* mice (Figure 6A). We used a KO-validated monoclonal antibody for ASC detection (Supplemental Figure 5A-Bb). 3D surface reconstruction of Iba1, GFAP and ASC immunosignal analysis were subsequently conducted (Figure 6A and Supplemental Figure 5C). We detected a significant increase in the total number of fluorescent ASC aggregates, especially in Iba1^+^ cells (likely microglia) at 7 dpi similar to the mRNA expression of *Asc* observed in the scRNA-Seq data analysis following mFPI (Figure 1B, Figure 6B and Supplemental Figure 5D). The number of ASC aggregates within Iba1^+^ cells followed the total ASC aggregates pattern (Figure 6B and Supplemental Figure 5C-D). Notably, *Nlrp3^−/−^*injured mice showed comparable ASC aggregates counts, both overall and within Iba1^+^ cells, compared to WT mice (Figure 6b) but significantly lower aggregates in GFAP^+^ cells (presumably reactive astrocytes, Figure 6c). Interestingly, at 21 dpi, the number of ASC aggregates outside Iba1^+^ and GFAP^+^ cells (extra-glial ASC) significantly increased in both WT and *Nlrp3^−/−^* mice (Figure 6d). However, this ensembles count was significantly lower in *Nlrp3^−/−^* mice compared to WT controls (Figure 6d). Additionally, as ASC aggregates numbers increased within microglia, their corresponding volumes also increased. To further analyze ASC aggregation, we categorized ASC aggregates by volume (1–20 µm^3^, 20–30 µm^3^, 30–40 µm^3^) and quantified the number of ensembles in each category (Supplemental Figure 5E-F and Figure 6E). ASC aggregates within a range of 30–40 μm^3^ were significantly decreased in the *Nlrp3^−/−^*mice at 21 dpi (Figure 6E). Collectively, our findings demonstrate a remarkable upregulation of ASC aggregates outside of Iba1^+^ and GFAP^+^ cells, along with marked ASC aggregation at 21 dpi. *Nlrp3^−/−^* mice displayed reduced ASC aggregation within Iba1^+^ cells and fewer extra-glia ASC aggregates compared to WT group at 21 dpi. These observations highlight the essential role of NLRP3 not only in promoting inflammasome activation after brain injury but also in facilitating significant ASC aggregation, as previously suggested in *in vitro* studies (41).

**Figure 6:**
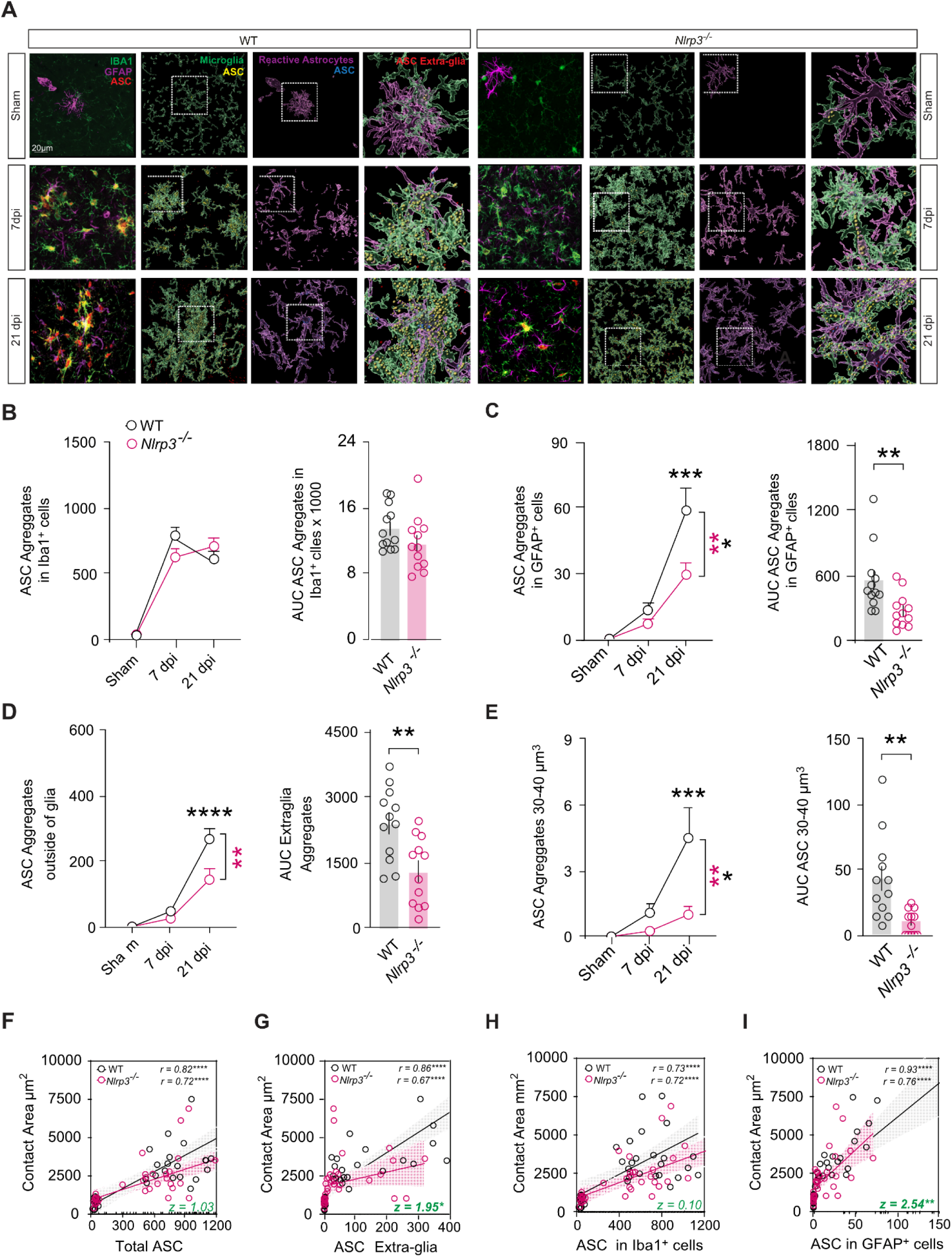
NLRP3 regulates ASC aggregation and distribution following CHI. **A.** Immunohistochemistry for Iba1 (green), GFAP (magenta) and ASC (red) in cortices of WT and *Nlrp3^−/−^*mice at 7 and 21 dpi. ASC localization within Iba1+ cells (yellow spots), GFAP^+^ cells (blue spots), and outside glial cells (red spots) were analyzed using IMARIS. Rectangles highlight examples of ASC distribution shown in high magnification views. Scale bar, 20 μm. **B-E.** Quantification of total ASC aggregates numbers (**B**), intracellular ASC aggregates within Iba1^+^ cells (**C**), ASC aggregates outside glia cells (**D**), and ASC aggregates with a volume range 30–40 um^3^ (**E**) at 7 and 21 dpi in WT and *Nlrp3^−/−^* mice subjected to Sham or CHI. *n* = 12 slices from 4 mice per group per each time point, \**p* < 0.05, \*\*\**p* < 0.01, \*\*\**p* < 0.001, \*\*\*\**p* < 0.0001 by ordinary two-way ANOVA with Bonferroni’s post hoc test. Right panels show integrative AUC of each marker over time (dpi). \**p* < 0.05, \*\**p* < 0.01, \*\*\**p* < 0.001, \*\*\*\**p* < 0.0001 by Student’s t-test. **F-I.** Spearman’s correlation analysis (*r*) with a linear regression model (black lines WT, red lines *Nlrp3^−/−^*) with an interaction term for ASC aggregates (total, within glial cells, and outside glial cells) and glia cell contact areas. n = 36 slice from 12 mice per genotype, \**p* < 0.05, \*\**p* < 0.01, \*\*\**p* < 0.001, \*\*\*\**p* < 0.0001. Differences between WT and *Nlrp3^−/−^* Spearman Correlation Coefficients (*r*) was performed using the standard Fisher’s z-transformation and subsequent comparison(69) \**p* < 0.05, \*\**p* < 0.01.

### Interaction between reactive astrocytes and microglia correlates with ASC distribution following CHI

To further investigate the relationship between reactive astrocytes and microglia interactions and ASC distribution following CHI, we conducted correlation analysis between ASC aggregates and glial contact areas after 3D reconstruction (Figure 6 F–I). Notably, we observed a strong positive correlation between the cell contact area and the number of extra-glia ASC aggregates *(r = 0.86, p < 0.0001)* (Figure 6g) as well as ASC aggregates within GFAP^+^ cells *(r = 0.93, p < 0.0001)* (Figure 6I), in WT animals after CHI. Although lower correlations coefficients were found between the reactive astrocytes and microglia contact areas and the number of ASC aggregates in microglia *(r = 0.73, p < 0.0001)* (Figure 6h) and the overall count of ASC aggregates *(r = 0.82, p < 0.0001)* (Figure 6f), these correlations remained significantly positive (Figure 6F–I). Interestingly, the genetic deficiency of the *Nlrp3* gene significantly reduced the correlation between the contact area and the number of extra-glia ASC aggregates *(z = 1.95, p < 0.05)* (Figure 6G) as well as ASC in GFAP^+^ cells *(z = 2.54, p < 0.01)* (Figure 6I). In summary, these results indicate that ASC aggregation after CHI is partly dependent on NLRP3 and additionally reactive astrocytes and microglia contacts are influenced by ASC expression, distribution, and aggregation, especially the ASC aggregates located outside of Iba1^+^ and GFAP^+^ cells.

## Discussion

In this study, we identify *Asc* as a key gen expressed in monocytes, macrophages, and especially in microglia populations during the subacute phase of CHI based on scRNA-seq data from mouse cortical cells after mFPI. Similarly, in our mild CHI mouse model, sustained ASC protein expression was confirmed, driving reactivity of microglia and astrocytes, and innate immune responses that contribute to neurobehavioral deficits following mTBI. Our findings not only complement and expand previous preclinical and clinical studies on moderate to severe TBI models (42, 43), which highlight the role of inflammasome cascades and their critical involvement in regulating secondary inflammatory responses after TBI, but also demonstrate for the first time that the main adaptor protein ASC is a key element in neuroinflammatory responses and cognitive impairment following mTBI. In contrast to several studies on moderate-to-severe TBI models, where the expression of inflammasome-related proteins typically peaks within the initial week following injury and declines over time (21, 44, 45), we found a progressive and sustained increase in inflammasome-related proteins specially after the first 7 days following mild CHI. This temporal pattern aligns with our observations of sustained morphological changes of reactive astrocytes and microglia during a later phase (21 dpi). These discrepancies between models and TBI types suggest that inflammasome engagement in innate immune responses and tissue repair varies depending on injury severity. Moreover, the prolonged pro-inflammatory characteristics exhibited by activated microglia and reactive astrocytes indicate that both glial cell types are capable of propagating inflammation through inflammasome-related pathways post-injury. Interestingly, our investigation demonstrated that astrocytes display more significant reactive morphology at 21 dpi compared to 7 dpi. This may result from the potential further induction of neurotoxic astrocytes by microglial inflammasome in response to sustained tissue damage, as previously reported in stress models (46). Furthermore, these findings align with existing evidence suggesting that distinct glial cell types respond differentially to brain injury in a temporal sequence (37, 47–49).

Additionally, we observed a widespread upregulation of inflammasome components following CHI, particularly at 21 dpi, with a significant increase in the cleaved forms of CASP1, CASP8 and IL-1β. Our results suggest the involvement of several ASC-dependent inflammasomes in the inflammatory pathways after CHI. This is evident by the pronounced reduction in CASP1, CASP3 and CASP8 levels and morphological changes of microglia and reactive astrocytes observed in *Asc^−/−^*mice. Most notably, our behavioral evaluation during the late subacute phases shows that anxiety-related behaviors (thigmotaxis), mild procedural learning deficits and long-term memory impairments were significantly diminished in animals with genetic deficiency on the *Asc* gene. Furthermore, our analysis established a significant positive correlation between reactive astrocyte and activated microglia interaction and the number of ASC aggregates. Notably, a potential connection has been identified between extracellular ASC and cellular propagation of inflammatory mediators, including IL-1β and TNF-α following inflammasome activation (50, 51). These cytokines not only promote the aggregation of glia and infiltrated cells around the trauma epicenter (52, 53), but may also influence cell communication far from the lesion site through extracellular ASC. ASC possess prion-like properties (51) and could propagate to neighboring glial cells, thereby triggering the activation of the ASC-mediated inflammasome cascade as previously shown in the context of Alzheimer’s disease-related amyloidosis (54, 55). These observations suggest a pivotal role of ASC in sustaining neuroinflammation and glial reactivity following CHI.

Our findings, however, contrast with a previous report on moderate CCI, where *Asc^−/−^* mice showed no differences in motor recovery or volume lesions (56). This inconsistency may suggest that, in the context of low-grade chronic neuroinflammation induced by mild, uncontrolled stimuli, such as a single concussion event, the expression, oligomerization, and propagation of ASC may occur gradually compared to the high-grade neuroinflammation triggered by severe injuries. Similarly, ASC-dependent inflammasome activation and pathological protein aggregation have been observed in several neurodegenerative disorders characterized by sustained chronic low-grade neuroinflammation (57), such as Alzheimer’s and Parkinson’s disease, where ASC plays a decisive role in the aggregating and further propagating Amyloid β (54) and α-Synuclein (58). Considering our findings, ASC may govern the immune molecular mechanism that triggers neurodegeneration and increase the risk of developing neurodegenerative conditions following TBI (59). Nonetheless, future investigations are still required to establish a comprehensive understanding of the specific temporal participation of inflammasomes across varying injuries severity and the risk factor for neurodegeneration.

There are certain limitations that should be acknowledged in our study. Focusing on the expression of inflammasome mediators and cell morphological characteristics may not fully capture the complex interactions with other compensatory mechanisms within the affected neuronal networks or the further anti-inflammatory modulation of secondary immune responses. Additionally, although our scRNA-Seq analysis determined specific cellular clusters upregulating *Asc* in the subacute phase of a CHI model, our study using conventional knockout line cannot dissect precisely the role of different peripheral and cellular actors in the immunological process following CHI. Thus, there is an imperative need to investigate the neuroinflammatory evolution of the specific neural, glial and peripheral cellular subtracts over extended post-traumatic periods.

Our findings pave the way to delve into specific pharmacological agents that can effectively modulate ASC expression, activity and aggregation and assess their impact on neurological outcomes after TBI. Considering our findings, pharmacological interventions targeting ASC may help to mitigate neuroinflammation and potentially promote neuroprotection, enhancing recovery of injury and preventing further neurodegeneration and disability.

## Methods

### Ethical compliance

We followed the European guidelines for animal research, conformed to the requirements of the German Animal Welfare Act and received approval from North Rhine-Westphalia State Agency for Nature, Environment and Consumer Protection under folder number 81-02.04.2019.A026 and 2024-791-Grundantrag.

### Single-cell RNA-sequencing data analysis

Raw FASTQ files and count matrices are publicly available in the NCBI GEO database under accession number GSE160763 (26). Single-cell RNA-seq data were analyzed using Seurat v5.1.0 in R v4.3.2 (60). Briefly, cells with over 5000 detected genes, UMI counts under 12,500, and less than 1% mitochondrial gene content were retained, followed by log-transformation of the data. We selected the top 4000 highly variable genes for principal component analysis (PCA) and used the top 20 principal components as input for UMAP dimensionality reduction. Cell cycle states were determined using Seurat’s CellCycleScoring (61) function with marker genes from Tirosh *et al.* (60). Batch correction was performed using Harmony to integrate datasets across samples (62). Clusters were identified at a resolution of 0.5. Cellular identities were assigned via automated cell type annotation using SingleR (29), which assigns cellular identity for single cell transcriptomes by comparison to reference data sets of pure cell types sequenced by microarray or RNA-sequencing (RNA-seq). Annotation relied on a reference dataset of 358 mouse RNA-seq samples annotated to 18 major cell types (63). Figures were generated using Tidyverse(64), SCpubr(65), and Enhanced Volcano (66).

### Mice

In this study, we used male and female mice aged 6–7 months with a C57BL/6 mixed genetic background (C57BL/6N-J) from our own animal facility (House of the Experimental Medicine) at the University Hospital Bonn in German. Wild-type (WT), *Nlrp3^−/−^*, and *Asc^−/−^* mice were initially sourced from Millennium Pharmaceuticals as previously published (67). All mice were backcrossed with C57BL/6N mice and maintained under pathogen-free, standardized conditions with a 12:12-hour light/dark cycle and ad libitum access to food and water. Behavioral experiments were conducted during the dark phase of the light/dark cycle.

### Electromagnetic controlled closed-head injury model of mild traumatic brain injury

To model mild TBI in mice, we adapted the electromagnetic controlled Closed-Head Model previously described (31) (Supplemental Figure 2A). Briefly, mice were anesthetized with 5% isoflurane during induction, and 1.5-2% isoflurane mixed with 100% oxygen (0.5-1 l/min) during surgery and fixed in a stereotaxic frame (Stoelting, Dublin, Ireland). A 1-mL latex pipette bulb filled with water was placed under the head to distribute the impact force. A midline sagittal scalp incision was made, and a single controlled midline skull impact was delivered at coordinates 0.0 mm mediolateral and -1.5 mm anteroposterior, with a velocity of 5 m/s, dwell time of 100 ms, and impact depth of 1.0 mm, using a stereotaxic electromagnetic impactor with a 5.0-mm steel tip (Stereotaxic Impactor, Leica Biosystems, Germany). Sham mice underwent identical surgical procedures without impact injury. In the post-operative phase, mice received carprofen (5 mg/kg) subcutaneously once daily and tramadol (1 mg/mL) in drinking water for three days.

### Morris water maze (MWM)

The evaluation of spatial learning and memory was conducted between 10 and 20 dpi. A circular pool with a diameter of 1 m was filled with white opacified water maintained at a temperature of 21–23 °C. The pool was dimly lit (approximately 40 lx) and surrounded by a white curtain. Asymmetrically placed distal cues were positioned on the pool wall to serve as spatial references. The pool was virtually divided into four quadrants, one of which contained a hidden platform (15 cm in diameter) submerged 1.5 cm below the water surface. Mice were trained to locate the platform using the distal cues for orientation. Training consisted of four trials per day for five consecutive days. During each trial, mice were placed into the water facing the pool wall in a quasi-randomized order to prevent them from developing fixed strategies. Mice were given 60 s to find the platform; if they failed to locate it within this time, they were manually guided to it. Once on the platform, mice were allowed to remain there for 15 s before the next trial began. After completing all four trials for the day, mice were dried and returned to their home cages. The integrated time or distance traveled during these trials was analyzed. A spatial probe trial was conducted 24 hours after the final training session (16 dpi). For this test, the hidden platform was removed, and mice were allowed to swim freely for 60 s. The drop position was in the quadrant opposite where the platform had been located, with each mouse facing the wall at the center of that quadrant. One day after the spatial probe trial, a visually cued learning phase began. During this phase, the platform was flagged for visibility and relocated to a different quadrant, while all distal cues were removed. Mice underwent three trials per day for three consecutive days (17–20 dpi) to learn to locate the flagged platform. All mouse movements were recorded and tracked using Noldus EthoVision software (Wageningen, Netherlands).

### Tissue preparation

Mice were deeply anesthetized using ketamine (100 mg/kg) and xylazine (20 mg/kg) and transcardially perfused with at least 30 ml of ice-cold PBS at designated time points (Sham, 1, 7, and 21 dpi). Following perfusion, brains were carefully dissected. One hemisphere was snap-frozen in liquid nitrogen and stored at -80°C until biochemical analyses, while the other hemisphere was fixed in 4% paraformaldehyde overnight. Coronal brain sections (40 μm thick) were obtained using a Leica VT1000S vibratome (Leica Microsystems Inc.) from the region spanning bregma -0.45 to -1.85 mm. Sections were collected in PBS at intervals of 300 μm for various staining procedures.

### Primary antibodies

For immunoblotting, following primary antibodies were used: Anti-Caspase 1 from AdipoGen (USA, mAb Casper-1, mouse anti-mouse, 1:1000), anti-ASC from Cell Signaling (Frankfurt, Germany, D2W8U, rabbit anti-mouse, 1:1500), anti-NLRP3 from Cell Signaling (Frankfurt, Germany, D4D8T, rabbit anti-mouse, 1:500), anti-IL-1β from Cell Signaling (Frankfurt, Germany, E7V2A, rabbit anti-mouse, 1:1000), anti-Caspase 3 from Cell Signaling (Frankfurt, Germany, D3E9, rabbit anti-mouse, 1:1000), and anti-Caspase 8 from Cell Signaling (Frankfurt, Germany, D5B2, rabbit anti-mouse, 1:1000). For immunohistochemical analysis, the antibodies included anti-Iba1 from Wako Chemicals (Neuss, Germany, rabbit anti-mouse, 1:1000), anti-IBA1 from Abcam (Cambridge, UK, ab289874, goat anti-mouse, 1:1000), anti-GFAP from ThermoFisher (Waltham, USA, 2.2B10, rat anti-mouse, 1:300), anti-IL-1β from R&D Systems (Minneapolis, USA, AF-401-SP, goat anti-mouse, 1:500), and anti-P53 from Abcam (Cambridge, UK, ab131442, rabbit anti-mouse, 1:500).

### Tissue protein extraction and immunoblotting

After thawing, brain tissue was homogenized in PBS containing 1 mM EDTA, 1 mM EGTA, and protease inhibitor cocktail Halt™ (Thermo Scientific, Waltham, Massachusetts, U.S.). The homogenates were extracted in RIPA buffer [25 mM Tris–HCl (pH 7.5), 150 mM NaCl, 1% Nonidet P-40, 0.5% NaDOC, 0.1% SDS] and centrifuged at 20000 ×g for 30 minutes. RIPA fractions were separated using NuPAGE® electrophoresis gels (Thermo Scientific, Waltham, Massachusetts, USA) and immunoblotted with primary antibodies, followed by incubation with the corresponding secondary antibodies. Immunoreactivity was detected with an Odyssey CLx Imaging System (LI-COR, Bad Homburg, Germany), and images were analyzed using ImageJ (NIH, Bethesda, USA).

### ELISA pro-inflammatory cytokine panel

The levels of IL-1β and TNF-α in RIPA fractions were measured using the V-PLEX Plus Mouse Pro-Inflammatory Panel 1 (Meso Scale Discovery, Rockville, USA), following the manufacturer’s protocol. Samples were diluted 1:1 on the plate with the reagent diluent provided in the kit. Signals were detected using a QuickPlex SQ 120 plate reader (Meso Scale Discovery, Rockville, USA).

### Immunohistochemistry

Brain sections were washed three times for 5 min with PBS and incubated in citrate buffer for 5 min at 95°C. After antigen retrieval, the sections were cooled down at room temperature and washed with PBS and in 0,5% Triton X-100 B (PBS-T), then blocked for 1 h with BSA 1% in PBS-T and incubated overnight with the primary antibodies. Next, the sections were washed three times for 5 min in PBS-T, incubated with appropriate secondary antibodies (1:1000) for 60 min, and following with three times with PBS washing for 5 min. The tissue was mounted using ProLong™ Gold Antifade Mountant with DNA Stain DAPI (Thermo Scientific, Waltham, Massachusetts, U.S.). Three coronal brain sections per animal (between bregma −1.00 and −1.85) were selected. Confocal z-stack images of the contusion were primarily acquired from layers 1 and 2/3 of the retrosplenial and primary motor cortex. Each z-stack consisted of 10 optical slices (c × z × t = 10), with one frame captured per acquisition. Imaging parameters were standardized across all samples: the pixel size was set to 0.1559814 µm in both the x and y axes, and the voxel depth (z-step) was maintained at 2.0 µm.

### Skeleton analysis of microglia and reactive astrocytes

Morphological changes in microglia and reactive astrocytes were quantitatively analyzed using a skeleton analysis method (68). Images were imported into ImageJ software (Bethesda, Maryland, USA), converted to grayscale, and adjusted for brightness and contrast to enhance cell process visibility. After obtaining binary images of microglia and reactive astrocytes, we used the plugin skeletonize to obtain skeletons, representing the medial axis of cell processes. Morphological parameters measured included: Process Branches, defined as number of branching points. Endpoints, defined as number of terminal points. Process Lengths, defined as total length of all processes. The data were normalized by dividing the measurements by the number of cells in each image.

### Contact analysis between microglia and reactive astrocytes

Imaris Image Analysis Software (version 9.5, Oxford Instruments, Oxon, England) was used to analyze the contact area between microglial and astrocytic surfaces. 3D reconstructions were created using the tool surfaces by setting intensity thresholds to differentiate cell surfaces from the background. Adjustments were made to ensure accurate cell boundary representation. The Surface-Surface Contact X-Tension identified and analyzed overlapping regions between microglial and astrocytic surfaces, generating a new surface to represent the contact area. The contact area (µm^2^) was calculated based on the number of overlapping voxels within the 3D reconstructions.

### Analysis of ASC aggregates

The cortical impact area was examined using a confocal LSM900 microscope (Zeiss, Oberkochen, Baden-Württemberg, Germany). Images were captured under consistent confocal settings and processed with Imaris Image Analysis Software (Oxford Instruments, Oxon, England) to identify ASC aggregates. Standardized settings were maintained across samples, including pinhole size, laser intensity, digital gain, and offset. Z-stacks were collected to obtain the complete 3D structure of cells and ASC aggregates. Image preprocessing included adjusting brightness and contrast, reducing noise, and subtracting background to enhance aggregate visibility. Imaris Image Analysis Software was used to create 3D reconstructions of microglia, astrocytes, and ASC aggregates. The “Surfaces” tool generated accurate 3D models by setting intensity thresholds. The Spots tool identified and quantified ASC aggregates based on fluorescence intensity and size, with thresholds set using positive (mCherry-ASC mice) and negative (*Asc^−/−^* mice) controls. The Spots Close to Surface tool analyzed the spatial distribution of ASC aggregates relative to microglia or astrocytes, indicating potential inflammasome activation.

*IL-1*β *analysis*

Co-immunostainings for IL-1β, Iba1 and GFAP were performed at 7 and 21 dpi. Three-dimensional reconstructions of microglia, reactive astrocytes, neuronal nuclei, and IL-1β signals were generated using Imaris Image Analysis Software (Oxford Instruments, Oxon, UK). The “Spots” tool quantified IL-1β signals in the 3D space by analyzing fluorescence intensity and particle size thresholds. To assess spatial relationships, the “Spots Close to Surface” tool measured the proximity of IL-1β signals to cellular surfaces, evaluating potential associations with microglia, astrocytes, or neurons. Quantitative metrics, including the number of IL-1β spots adjacent to specific cell types, were measured for statistical analysis.

### Statistics and reproducibility

Data other than scRNA-seq were analyzed using GraphPad Prism Software version 9.0 or later (GraphPad Software, Inc., La Jolla, CA, USA) and are presented as mean ± SEM. After assessing the distribution, statistical comparisons were performed using either a two-tailed t-test or one- or two-way ANOVA with Bonferroni’s or Tukey’s post hoc test to identify group differences. Morris Water Maze (MWM) learning data were analyzed using repeated-measures two-way ANOVA with Tukey’s post hoc test. Correlation analyses were performed using non-parametric Spearman correlations. Differences between Spearman correlation coefficients were assessed using the standard Fisher’s z-transformation and subsequent comparison(69). Results were considered statistically significant at p < 0.05.

### Data availability

The data that supports the findings of this study are available on request from the corresponding authors upon reasonable request.

### Code availability

*p*Code used in analyzing scRNA-seq data is available at https://gitlab.lcsb.uni.lu/pablo.botellalucena/rna_single_seq.

## Author contributions

Conceptualization: SCG, MTH. Methodology: ASV, PBL, SCG, TL, SS, YD. Investigation: ASV, PBL, SCG, TL, SS, YD. Funding acquisition: SCG, MTH. Supervision: SCG, MTH. Writing of original draft: SCG, TL, YD. Writing, review, and editing: MTH, SCG, PB, TL, YD, YD, VS, EL

## Acknowledgements

We thank Ildikó Rácz and Christina Ising for their help in obtaining approval from the local ethics committee for the animal experiments; Sean-Patrick Hermann for assistance with editing the final draft; Juan I. Muñoz, Paula Martorell, and Nadia Villacampa for valuable discussions; and the DZNE Light Microscopy Facility for support with imaging and analysis. PBL is supported by the Fonds National de la Recherche within the PEARL program (FNR/16745220). SCG is supported by the Alzheimer Forschung Initiative e.V (Grant 21060), the Hertie Network of Excellence in Clinical Neuroscience (Grant 2021-1A-12), the Bonn Neuroscience Clinician Scientist Program (Neuro-aCSis), and the DFG (EXC2151– 390873048). TL was supported by Department of Biotherapy Cancer Center and State Key Laboratory of Biotherapy, West China Hospital, Sichuan University, Chengdu, China. Y.D. is supported by China Scholarship Council (CSC, File No. 202308310126). Y.D. is supported by China Scholarship Council (CSC, File No. 201908510162).

---

**Supplemental Figure 1:**
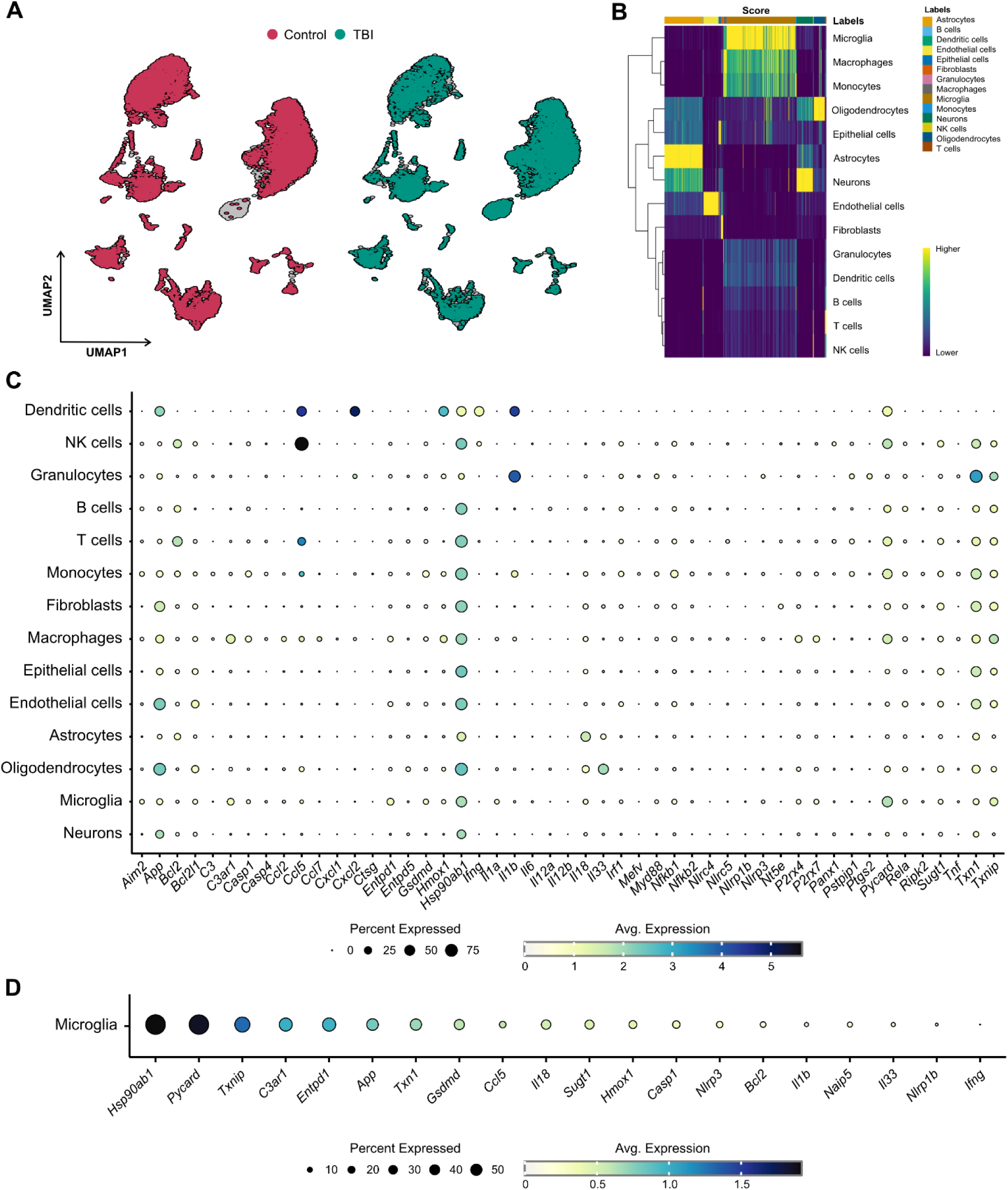
Single-Cell Transcriptomic Analysis of Brain Immune Cells Following TBI. **A.** UMAP clustering of single-cell RNA sequencing (scRNA-seq) data, comparing control (red) and TBI (green) samples, illustrating shifts in immune cell populations post-TBI. **B.** Heatmap showing gene expression profiles across different brain cell types and cluster identification. **C.** Dot plot depicting the expression of key inflammasome-related genes across different brain cell types. **D.** Dot plot depicting gene expression profile of inflammasome-related genes in microglia after TBI.

**Supplemental Figure 2:**
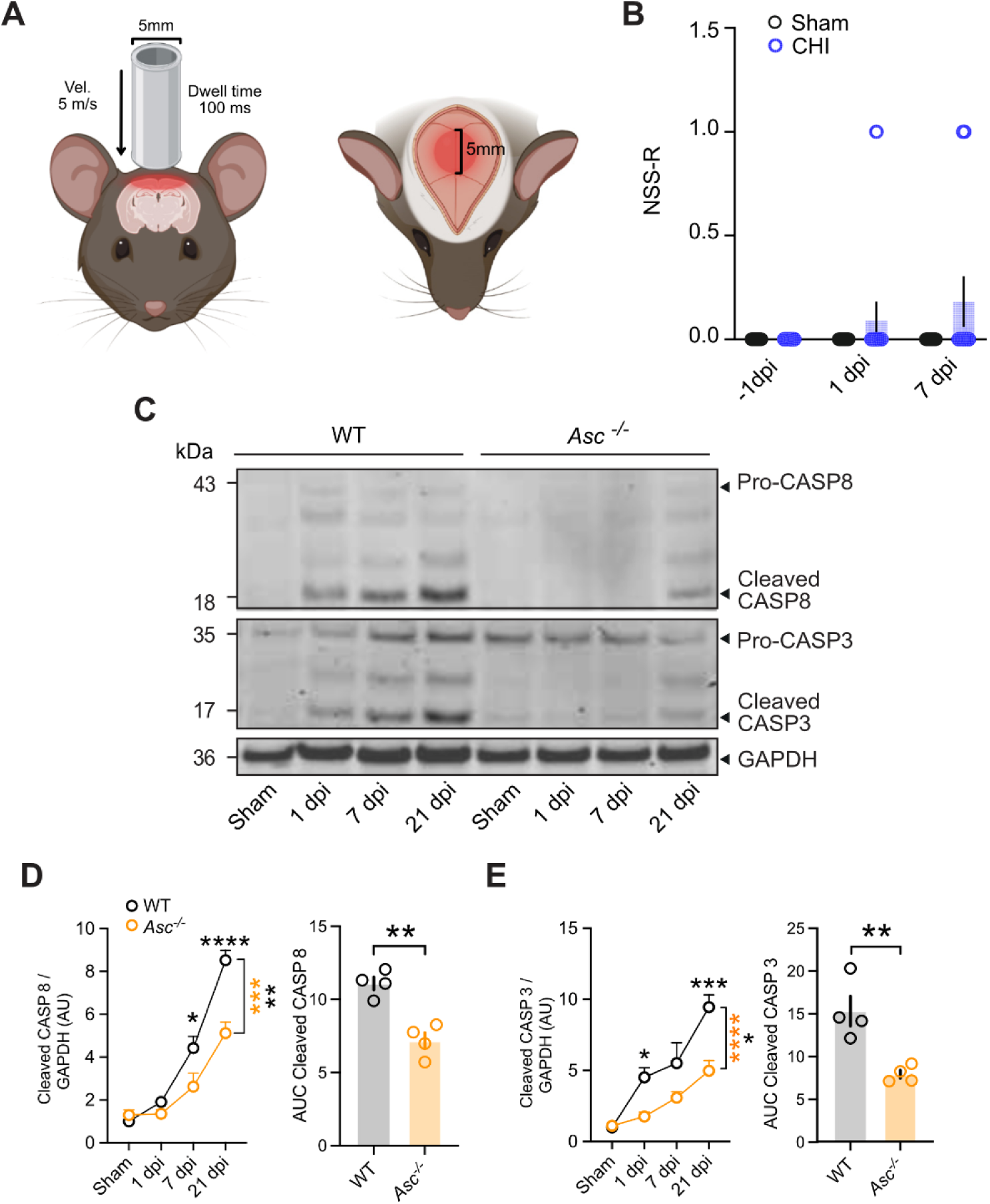
ASC modulates expression and cleavage of CASP-8 and CASP-3 following CHI. **A.** Schematic of injury model/location. Coronal view (right) shows tip positioned over the skull midline. Horizontal view (left) shows impact site relative to bregma. **B.** No significant motor, reflex or reaction deficits at 1- or 7-days post-intervention (dpi) when the mice were evaluated using the Revised Neurobehavioral Severity Scale (NSS-R). (Two-way ANOVA followed by Bonferroni’s post-hoc tests, n=11 mice per group per each time point, data are presented as the mean ± SEM) **c.** Representative immunoblot images of CAPS 8 and CASP 3 at 1, 7, and 21 dpi in peri-contusional cortices of mice subjected to Sham or CHI, comparing WT and *Asc ^−/−^* groups. **D-e.** Quantification of expressions of CASP 8 and CASP 3. The band intensity of a given target protein was normalized to the corresponding GAPDH signal for each sample. Data were further normalized to the average of the corresponding WT group and are presented as the mean ± SEM. Statistical significance was determined using two-way ANOVA followed by Bonferroni’s post-hoc tests, n = 4 mice per group per each time point, *p < 0.05, **p < 0.01, ***p < 0.001, ****p < 0.0001 (left panels). Quantification of area under the curve (AUC) of each target over time (dpi) in WT and Asc*^−/−^*mice is shown in right panels. Statistical analysis was performed using an unpaired two-tailed t-test. Data are presented as the mean ± SEM. *p < 0.05, **p < 0.01, ***p < 0.001, ****p < 0.0001.

**Supplemental Figure 3:**
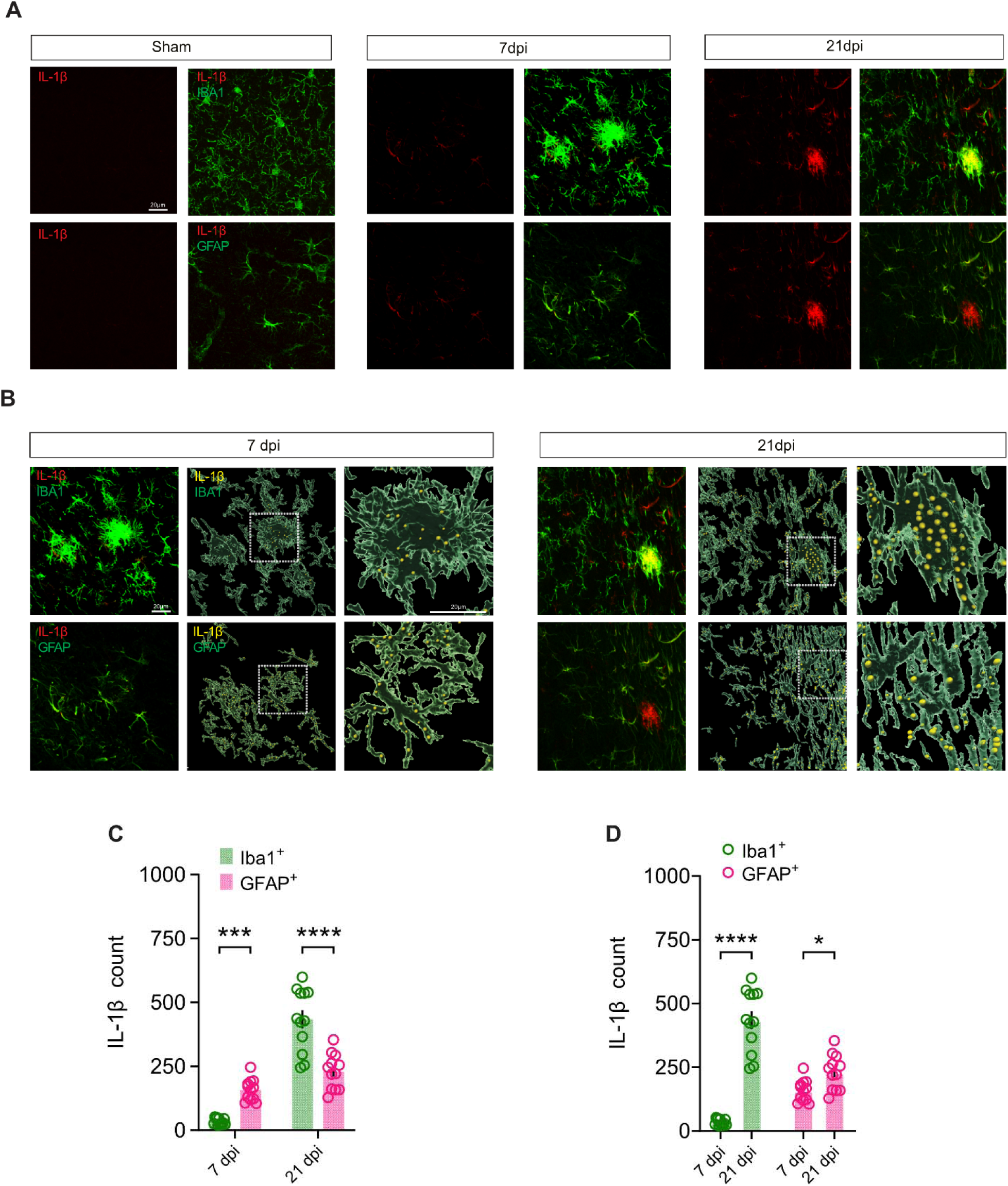
The cellular source of IL-1 β Following mTBI. **A.** Representative images of immunohistochemical staining of the contusional cortices for IL-1β (red), Iba1 (green) and GFAP (green) in mice following sham surgery and 7dpi and 21 dpi after CHI. **B.** 3D reconstruction of IL-1β immunostaining at 7dpi and 21 dpi following CHI. IL-1β (yellow spot) localization within Iba1^+^ and GFAP^+^ cells. Scale bar, 20 μm. **C-D.** IL-1β spot counts assessed in Iba1^+^ and GFAP^+^ cells at 7 and 21 dpi following CHI. Data are presented as the mean ± SEM. Statistical significance was determined using two-way ANOVA followed by Bonferroni’s post-hoc tests, n = 12 slice (4 mice) per group per each time point, *p < 0.05, **p < 0.01, ***p < 0.001, ****p < 0.0001.

**Supplemental Figure 4:**
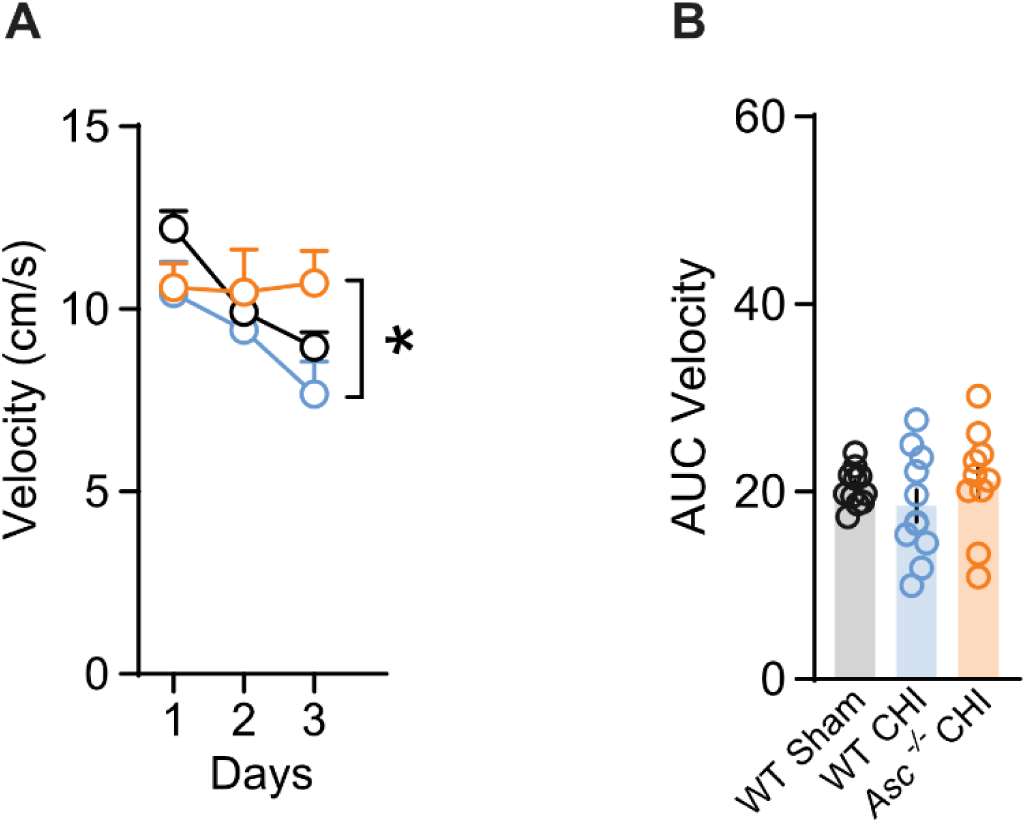
Swimming velocity during cued learning in the Morris water maze. Swimming velocity during cued learning in Sham WT, CHI-injured WT **A.** CHI-injured *Asc^−/−^*. **B.** Comparison of the integrated area under the curve of swimming velocity.

**Supplemental Figure 5:**
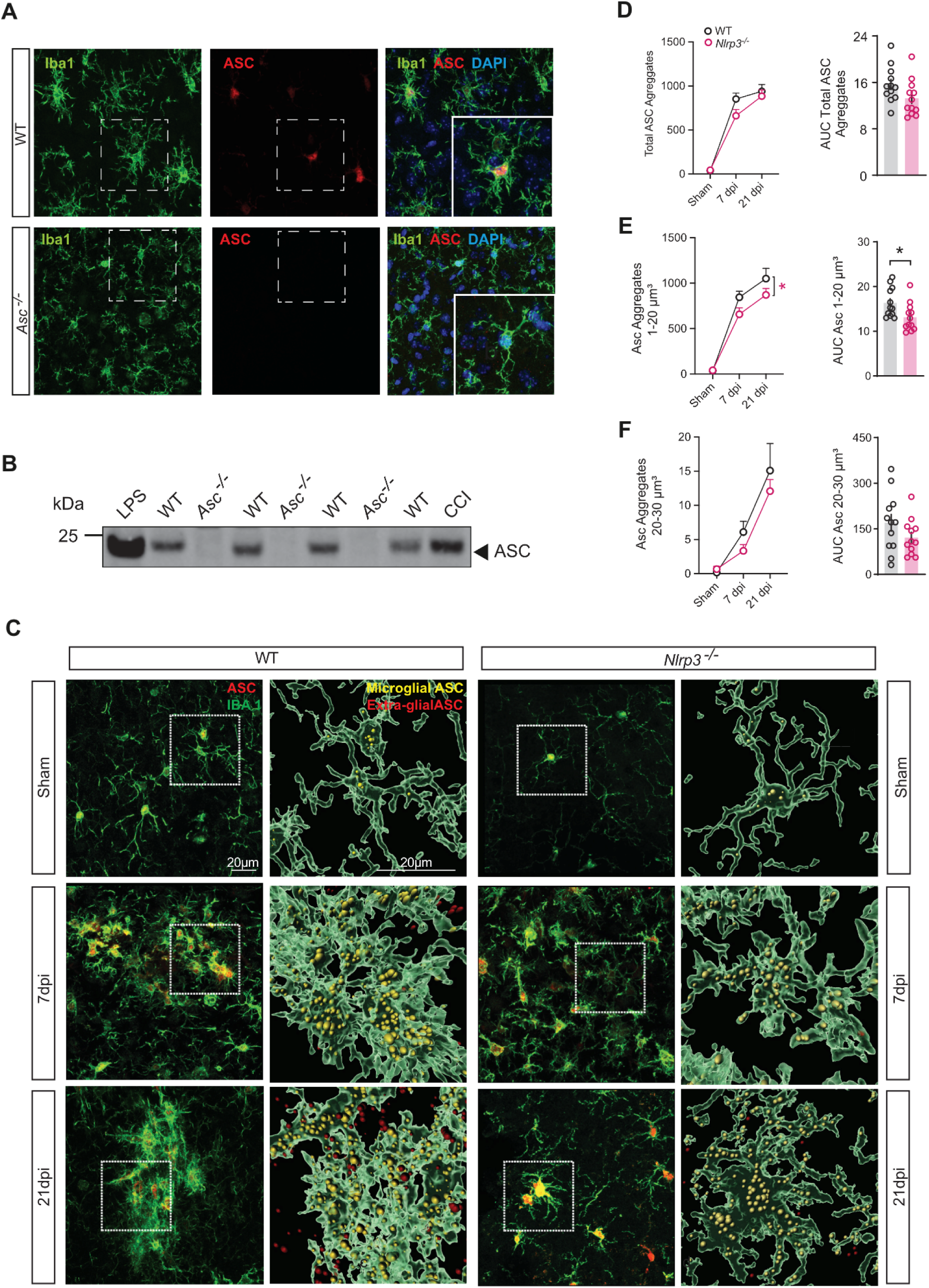
ASC antibody validation and ASC aggregates analysis. A-B. Validation of ASC antibody specificity using immunofluorescence staining and immunoblotting. **A.** ASC signal is detected in WT mice but absent in *Asc^−/−^* mice. **B.** Precipitated supernatant from LPS-stimulated primary microglia and cortical tissue from CCI brain-injured mice were used as positive controls. **c.** Representative immunohistochemical images of Iba1 (green) and ASC (red) in peri-contusional cortex of WT and *Nlrp3^−/−^* mice at different time points (sham, 7dpi, 21 dpi). ASC aggregates were analyzed and 3D reconstructed using IMARIS. Scale bar, 20 μm. **D-F.** Quantification of ASC aggregation and ASC aggregates volume. Two-way ANOVA with Bonferroni’s post-hoc tests (*p < 0.05, **p < 0.01, ***p < 0.001, ****p < 0.0001). n = 12 slices (4 mice) per group per each time point. Data shown as mean ± SEM.

